# A novel whole yeast-based subunit oral vaccine against Eimeria tenella in chickens

**DOI:** 10.1101/2021.11.05.467441

**Authors:** Francesca Soutter, Dirk Werling, Matthew Nolan, Tatiana Küster, Elizabeth Attree, Virginia Marugán-Hernández, Sungwon Kim, Fiona M. Tomley, Damer P. Blake

**Author notes:** **Correspondence:** Damer Blake and Dirk Werling.

## Abstract

Cheap, easy-to-produce oral vaccines are needed for control of coccidiosis in chickens to reduce the impact of this disease on welfare and economic performance. *Saccharomyces cerevisiae* yeast expressing three *Eimeria tenella* antigens were developed and delivered as heat-killed, freeze-dried whole yeast oral vaccines to chickens in four separate studies. After vaccination, *E. tenella* replication was reduced following low dose challenge (250 oocysts) in Hy-Line Brown layer chickens (p<0.01). Similarly, caecal lesion score was reduced in Hy-Line Brown layer chickens vaccinated using a mixture of S. cerevisiae expressing EtAMA1, EtIMP1 and EtMIC3 following pathogenic-level challenge (4,000 *E. tenella* oocysts; p<0.01). Mean body weight gain post-challenge with 15,000 *E. tenella* oocysts was significantly increased in vaccinated Cobb500 broiler chickens compared to mock-vaccinated controls (p<0.01). Thus, inactivated recombinant yeast vaccines offer cost-effective and scalable opportunities for control of coccidiosis, with relevance to broiler production and chickens reared in low-and middle-income countries (LMICs).

## 1 Introduction

Coccidiosis, a disease of the gastrointestinal tract caused by *Eimeria* parasites, is a considerable burden to the poultry industry economically, estimated to cost over £10 billion per year (1), and in terms of chicken health and welfare, causing diarrhoea and ill-thrift. Existing vaccines that consist of controlled oral doses of live or live-attenuated parasites are efficacious and widely used in egg-laying and breeder chicken populations. However, despite achieving levels of protection comparable to anticoccidial drugs (2, 3), uptake in the broiler chicken sector has been limited, at least in part because the cost of vaccines is relatively high (1). Furthermore, the current live anticoccidial vaccines can only be produced by controlled infection of specific-pathogen-free chickens, creating an inherent limit on productive capacity and questions around the ethical use of chickens for production of vaccines. Even if demand for existing vaccines increases, it is unlikely that the production of live-attenuated vaccines could be scaled up sufficiently to supply the entire broiler sector. Thus, novel oral vaccines against coccidiosis in chickens would provide a much-needed alternative to these current vaccines as well as in-feed anticoccidial drugs.

Recombinant protein vaccines, both subunit and “live” recombinant, targeting *Eimeria* species have long been examined as an alternative to current vaccines, and a number of immunoprotective antigens, such as Apical Membrane Antigen 1 (AMA1), Immune Mapped Protein 1 (IMP1) and Microneme Protein 3 (MIC3) have been identified (reviewed in Blake et al., 2017). Reduced parasite replication and gut pathology have been demonstrated when these antigens were given as subunit vaccines such as *E.coli* expressed recombinant proteins, DNA vaccines or administered using vectored approaches such as expression in transgenic *Eimeria.* For example, proof-of-concept vaccination trials administering three sub-cutaneous or intramuscular doses of recombinant protein have been successful at reducing parasite replication (faecal oocyst count) following low dose parasite oral challenge (250 oocysts) (4–6). However, the delivery of recombinant vaccines by intramuscular injection or other individual bird-by-bird approaches is unsuitable for large scale vaccination of chickens in the field, inhibiting commercialisation. Oral recombinant subunit vaccines that stably express candidate immunoprotective antigens provide an easier method of delivery to chickens in all environments and are therefore highly desirable.

Yeasts, such as *Saccharomyces cerevisiae*, have long been utilized for the production of recombinant soluble proteins for use in applications such as vaccination(7). More recently, *S. cerevisiae* has been used for oral vaccine delivery as whole recombinant yeast (live or killed), combining multiple purposes such as antigen expression and carriage, as well as being its own adjuvant. In addition to convenience for vaccine delivery, oral administration of recombinant S. cerevisiae supports direct vaccine interaction with the mucosal surface of the gastrointestinal tract, where *Eimeria* parasites invade, and can stimulate an immune response through pattern recognition receptors (PRRs), negating the need for adjuvants (8). Furthermore, *S. cerevisiae* and its cell wall derivatives are already used as feed supplements in the poultry sector and some studies have demonstrated its efficacy in reducing reproduction of *Eimeria* parasites (9) as well as improved production parameters during infection (10, 11), even without the addition of a vaccinal antigen. Previous studies of *S. cerevisiae* expressing microneme proteins have demonstrated efficacy in reducing oocyst output and caecal lesions following vaccination with live oral yeast (12, 13). Finally, recombinant yeast lines can be inactivated, permitting Generally Regarded as Safe (GRAS) status that may improve industry and social acceptance (14).Furthermore it has been suggested that inactivation could enhance antigenicity of yeast lines by altering distribution and exposure of structures such as β-1,3 glucan on the yeast cell surface (15).

The aim of this study was to produce a whole yeast vaccine using recombinant *S. cerevisiae* expressing the *Eimeria tenella* antigens EtAMA1, EtIMP1 and EtMIC3, and to assess efficacy in reducing challenge-induced parasite replication and pathognomonic lesions in the caeca in vivo following yeast inactivation and oral vaccination. The efficacy of vaccination using yeast expressing single *E. tenella* antigens or combinations of yeast expressing each antigen are assessed and compared to E.coli expressed, purified recombinant protein and mock live oocyst vaccination following low or high dose parasite challenge.

## 2 Materials and Methods

### 2.1 Ethics statement

This study was performed under a UK Home Office License according to the Animals in Scientific Procedures Act 1986 (ASPA). Procedures were approved by the Royal Veterinary College (RVC) Animal Welfare Ethical Review Body (AWERB).

### 2.2 Vaccine antigens

Three *E. tenella* (Houghton strain) antigens were selected for surface expression on *S. cerevisiae*; the ectodomain of Apical Membrane Antigen 1 (EtAMA1), Immune Mapped Protein 1 (EtIMP1) and repeat 3 from Microneme Protein 3 (EtMIC3), one of three identical Microneme Adhesive Region (MAR) domains contained within the ectodomain (16). DNA sequences for EtAMA1, EtIMP1 and EtMIC3 were obtained from Genbank, (accession numbers LN609976.1, FN813229.2 and FJ374765.1 respectively). Codon optimisation for *S. cerevisiae* was performed using the codon usage database (http://www.kazusa.or.jp/codon) for EtAMA1 and EtIMP1, but synthesis was unsuccessful for EtMIC3.

### 2.3 Cloning strategies applied

#### Study 1

The *E. tenella* AMA1 ectodomain coding sequence, representing amino acids 24-446, was combined with a 3’codon optimised citrine tag (EtAMA1Cit) in the first study for assessment of protein expression. In parallel, the *E. tenella* IMP1 coding sequence (amino acids 2-387) was also combined with a 3’ citrine tag (EtIMP1Cit). DNA constructs were synthesised by Eurofins Genomics (Luxembourg) within a pEX-K4 (EtAMA1Cit) or pMK-RQ (EtIMP1Cit) plasmid with appropriate restriction enzyme sites for cloning into pYD1 yeast display plasmid vector (Invitrogen, Thermofisher Scientific, Waltham, MA, USA). A NotI restriction site was included between each antigen sequence and the 3’citrine tag. Attempts to synthesise the EtMIC3 R3 sequence were unsuccessful and thus this antigen was not included in Study 1.

Synthesised antigen constructs were transformed into competent XL-1 Blue *Escherichia coli* cells (Agilent Technologies, Santa Clara, CA, USA) by electroporation, according to manufacturer’s instructions and pipetted on to selective Lysogeny Broth (LB) agar plates (50 μg mL^−1^ kanamycin, 20 mM IPTG, 80 μg mL^−1^ X-gal, all Sigma Aldrich, St Louis, MO, USA) and incubated overnight at 37 °C. Blue/white colony screening was performed, and selected colonies grown in LB broth (50 μg mL^−1^ kanamycin) overnight at 37 °C with shaking at 160 rpm. Plasmid DNA was prepared from overnight cultures using the QIAprep Spin Miniprep kit (QIAGEN, Hilden, Germany) according to manufacturer’s instructions.

Restriction digest of pEX-K4 plasmid DNA containing EtAMA1Cit or was performed using *Bam* HI and *Xho* I (New England Biolabs, Ipswich, MA, USA) to extract tagged antigen coding sequences for cloning into pYD1 plasmid vector. Restriction digest of pYD1 plasmid was performed with the same restriction enzymes. Restriction digest of pMK-RQ plasmid DNA containing EtIMP1 was performed using *Bam* HI and *Not* I restriction enzymes (New England Biolabs). Restriction digest of pYD1-EtAMA1Cit plasmid was performed with the same restriction enzymes to remove the EtAMA1 insert but retain the citrine tag. Restriction digest products were visualised by gel electrophoresis on a 0.8 % agarose gel and bands cut out using a scalpel blade. Gel extraction was performed using the QIAquick gel extraction kit (QIAGEN) according to manufacturer’s instructions and concentration quantified by spectrophotometry using the DS-11 FX spectrophotometer (Denovix, Wilmington, DE, USA). Ligation was performed on gel purified insert and digested plasmid with T4 Ligase (Promega, Madison, WI, USA) according to manufacturer’s instructions, ligation reactions were transformed into XL-1 Blue *E. coli* cells (Agilent Technologies) by electroporation as before and pipetted on to selective LB agar plates (100 μg mL^−1^ Ampicillin, 20 mM IPTG, 80 μg mL^−1^ X-gal, all Sigma Aldrich). Blue/white colony screening was performed, and selected colonies sent for Sanger sequencing at Eurofins Genomics to confirm correct integration of EtAMA1Cit into the pYD1 plasmid or EtIMP1 into the pYD1-Cit plasmid. Following confirmation by sequencing, plasmid DNA was prepared from overnight cultures as before and then used for yeast transformation. Competent *S. cerevisiae* EBY100 strain cells (Thermofisher Scientific) were transformed using the S.c. EasyComp™ Transformation Kit (Thermofisher Scientific) according to manufacturer’s instructions. Transformants were grown on minimal dextrose plates supplemented with 1 % leucine and 2 % glucose at 30 °C for 3-5 days. Empty (undigested) pYD1 plasmid DNA was also transformed into *S. cerevisiae* EBY100 strain.

#### Studies 2,3 and 4

To optimize antigen expression in yeast, plasmids were re-constructed without the citrine tag. EtAMA1 and EtIMP1 coding sequences (codon-optimised for yeast) were excised from the constructs generated in study 1 by restriction digest with BamHI and NotI (New England Biolabs). Cloning of the untagged EtAMA1 and EtIMP1 coding sequences into pYD1 was then performed as described for study 1.

The target *E. tenella* MIC3 R3 (EtMIC3) sequence (not codon-optimised for yeast) had previously been cloned into pET22b plasmid (MerckMillipore, Burlington, MA, USA) and transformed into XL-1 Blue *E. coli* cells (Agilent Technologies) in another study and stored in glycerol at −80 °C (16). A sub-sample was streaked on to selective LB agar plates (100 μg mL^−1^ Ampicillin) and incubated overnight at 37 °C. Selected colonies grown in LB broth (100 μg mL^−1^ Ampicillin) overnight at 37 °C with shaking at 160 rpm. Plasmid DNA was prepared from overnight cultures using the QIAprep Spin Miniprep kit (QIAGEN) according to manufacturer’s instructions.

PCR was used to amplify EtMIC3 DNA from the pET22b MIC3 plasmid using primers (F: GCTATCGGATCCCAAGCCGTTCCAGAGG, R: CTGCGAGAATTCGCCACTTGGATCTTCCGTT, 0.4 μM final concentration, Sigma Aldrich) that incorporated appropriate restriction enzyme sites (*Bam* HI and *Eco* RI) for cloning into pYD1. Each 50 μL reaction contained 5 μL High Fidelity PCR Buffer 10x, 1 μL dNTP mix (0.2 mM final concentration each), 2 μL MgSO4 (2 mM final concentration) and 0.2 μL Invitrogen™ Platinum™ Taq DNA Polymerase High Fidelity(1IU; all ThermoFisher Scientific).

PCR was performed using a G-Storm GS1 Thermal Cycler (Gene Technologies). Reactions were heated to 94 °C for 1 min, followed by 35 cycles consisting of 94 °C for 15 s, 55 °C for 30 s and 68 °C for 1 min. PCR products were visualised by gel electrophoresis on a 0.8 % agarose gel and extracted from the gel as before. Restriction digest of EtMIC3 PCR product was performed using *Bam* HI and *Eco* RI restriction enzymes (New England Biolabs) to extract antigen coding sequence for cloning into pYD1 plasmid vector. Cloning of EtMIC3 into pYD1 was then performed as described for study 1.

Transformation of pYD1 plasmids containing EtAMA1, EtIMP1 and EtMIC3 into *S. cerevisiae* EBY100 yeast was carried out as described above for study 1.

### 2.4 Production of yeast and confirmation of expression

*Saccharomyces cerevisiae* EBY100 yeast transformed with pYD1 including *E. tenella* antigen coding sequences (Study1: EtAMA1Cit, EtIMP1Cit; Studies 2-4: EtAMA1, EtIMP1, EtMIC3) were grown on minimal dextrose plates (0.67 % Yeast Nitrogen Base (YNB), 1.5 % agar, both Sigma Aldrich) supplemented with 1 % leucine (Sigma Aldrich) and 2 % glucose (Sigma Aldrich) at 30 °C for 3-5 days. Single colonies were inoculated into YNB-CAA (0.67 % YNB, 0.5 % Casamino acids (CAA), Calbiochem, San Diego, CA, USA) medium containing 2 % glucose and grown overnight at 30 °C plus shaking 200 rpm in an orbital shaker (New Brunswick™ Excella® E24 Shaker). Overnight cultures were centrifuged at 4,000 × G for 10 min at room temperature and resuspended in YNB-CAA containing 2 % galactose (Sigma Aldrich) to an OD_600_ of 1.0 to induce protein expression. Yeast were cultured for 24 h at 30 °C plus shaking 200 rpm in an orbital shaker.

Confirmation of protein expression 24 h post-induction was assessed by antibody staining and flow cytometry. A volume of yeast equal to an OD600 of 2.0 was pelleted by centrifugation at 6000 × G for 3 min and washed in PBS. Cell pellets were then incubated with a mouse Anti-V5 tag monoclonal antibody (1.2 mg mL^−1^; Thermofisher Scientific) in PBS 0.1% BSA for 45 min at 4 °C. Cells were then washed twice in PBS then suspended in goat anti-mouse IgG (H&L) cross-adsorbed secondary antibody Alexa Fluor 488 (2 mg mL^−1^; Thermofisher Scientific) in PBS 0.1 % BSA for 45 min at 4 °C. Cells were then washed twice with PBS and then resuspended in FACSFlow (Becton Dickinson, Franklin Lakes, NJ, USA) prior to analysis using a FACSCalibur (Becton Dickinson). Expression was analysed using the FlowJo software package (V10, FlowJo LLC, Ashland, OR, USA), by comparing expression of the V5 tag, expressed at the 3’ end of the antigen coding sequence, at 24 h post-induction compared with the staining obtained prior to induction.

For study 1, yeast (24 h post-induction) were counted using the TC20™ automated cell counter (Bio-Rad Hercules, CA, USA). Cells were centrifuged at 4,000 × G for 10 min and resuspended in PBS (Thermofisher). Yeast cells were heat-treated at 56 °C for 1 h, pelleted and then freeze-dried overnight using a Lyodry compact (Mechatech Systems Ltd, Bristol, UK) and stored at −20 °C until oral inoculation into chickens. 1.7 × 10^7^ yeast cells were resuspended and delivered to each chicken by oral gavage in 100 μL of PBS. For chickens receiving both EtAMA1 and EtIMP1, 50 μL of each was combined and 1.7 × 10^7^ yeast cells in total were delivered to each chicken in 100 μL of PBS.

For studies 2, 3 and 4, yeast (24 h post-induction) were counted as described. Cells were centrifuged at 4,000 × G for 10 min and resuspended in PBS (Thermofisher) to a concentration of 1.5 × 10^7^ cells mL^−1^. 1 ml aliquots were heat-treated at 95 °C for 2 min, pelleted and then freeze-dried overnight as before. Freeze-dried yeast were stored at 4 °C and then resuspended in individual doses of 600 μL PBS 24 h prior to oral inoculation of yeast into chickens. For chickens receiving all three yeast expressing antigens, each was resuspended in 600 μL and then 200 μL of each yeast was mixed in one microcentrifuge tube for oral dosing.

Confirmation of successful yeast killing was confirmed by pipetting 50 μL of each heat-killed yeast (at two concentrations: 1.5 × 10^7^ cells mL^−1^ and 4.9 × 10^8^ cells mL^−1^) on to minimal dextrose plates supplemented with 1 % leucine and 2 % glucose at 30 °C. Killing was confirmed by the absence of growth after 5 days incubation. Second, ~1.5 × 10^7^ heat killed yeast cells were diluted in 5 ml YNB-CAA medium containing 2 % glucose and grown for five days at 30 °C plus shaking 200 rpm in an orbital shaker. Growth was assessed by spectrophotometry, comparing OD_600_ of heat killed and sterile (i.e. no yeast) broths.

### 2.5 Animals

For studies 1,2 and 3, female Hy-line Brown layer chickens were purchased at day of hatch from Hy-line UK Ltd (Studley, UK). All layer chickens were vaccinated against Marek’s disease (Nobilis Rismavac+CA126, MSD, Milton Keynes, UK) at the hatchery prior to the start of the study. Layer chickens were fed a commercial organic starter feed, free from anticoccidial drugs. For study 4, Cobb500 broiler chickens were purchased from P. D. Hook (Hatcheries) Ltd. (Cote, UK), at day of hatch. Cobb500 broiler chickens were vaccinated against infectious bronchitis (Nobilis IB H120, MSD Animal Health, Milton Keynes, UK). Broiler chickens were fed ad-lib throughout the study receiving starter feed from day 0-8, grower feed from day 9-18 and finisher feed from day 19 until the end of the study, all feeds were free from anticoccidial drugs (Target feeds, Whitchurch, Shropshire, UK) (Supplementary Table 1).

### 2.6 Parasites

The *E. tenella* Houghton (H) reference strain was used in this study (17). Parasites were passaged through chickens at the Royal Veterinary College as originally described by (18) and were used within three months of sporulation.

### 2.7 Experimental design

#### Study 1 (Low dose challenge-layer chickens)

A low parasite dose challenge study was used to assess vaccine efficacy against *E. tenella* replication, recognising that quantification of replication following higher doses can be complicated by the *Eimeria* crowding effect (19). Forty-eight female Hy-Line Brown layer day of hatch chicks were weighed and divided into eight groups of six to seven chicks, so that each group contained a mixture of chickens of approximately the same weight (Supplementary Table 2), each group was housed in a separate cage. At day 7 all birds were wing tagged for identification of individual chickens. Four groups received an oral yeast vaccine by oral gavage every 3-4 days from day 7 of age (five doses per chicken in total); empty vector (pYD1 only), pYD1-EtAMA1Cit, pYD1-EtIMP1Cit or an equal mixture of pYD1-EtAMA1Cit and pYD1-EtIMP1Cit. One group received a low dose live oocyst “vaccine” of 100 *E. tenella* oocysts by oral inoculation at day 7 of age(20), although vaccine recycling was much reduced by accommodation in wire floored cages preventing chicken access to most faecal material. One group received *E.coli* expressed recombinant EtIMP1 protein, prepared as previously described (6), by intramuscular injection at day 7 and day 15. Two groups did not receive any vaccination. All groups except one (unvaccinated, unchallenged) were challenged at day 22 with 250 sporulated *E. tenella* oocysts. All chickens were weighed and culled five days later. The left caeca were collected immediately and stored in RNAlater™ (Thermofisher Scientific) at 4 °C prior to homogenisation.

#### Studies 2 and 3 (High dose challenge-layer chickens)

High parasite dose challenge studies were used to assess vaccine efficacy against pathological (e.g. intestinal lesion scoring) and performance (e.g. body weight gain) parameters(21). In study 2, 100 female Hy-Line Brown layer day of hatch chicks were weighed and divided into eight groups of 12-13 chicks (Supplementary Table 2). Groups were housed in separate cages, with live oocyst vaccinated and unvaccinated/unchallenged groups isolated in separate rooms. In study 3, 210 female Hy-Line Brown layer day of hatch chicks were weighed and divided into six groups of 33-34 chicks, each group was housed in a separate rack of three cages with *E. tenella* challenged and unchallenged groups isolated in separate rooms (Supplementary Table 2). At day 7 all birds were wing-tagged. Both studies followed the same timetable except that in study 2, one group received a low dose live oocyst “vaccine” of 100 *E. tenella* oocysts by oral inoculation at day 7 of age.

Both studies received an oral yeast vaccine every 3-4 days from day 7 of age (five doses per chicken in total). In study 2, there were five groups receiving an oral yeast vaccine; empty vector (pYD1 only), pYD1-EtAMA1, pYD1-EtIMP1, pYD1-EtMIC3 or a mixture of pYD1-EtAMA1, pYD1-EtIMP1 and pYD1-EtMIC3 (Supplementary Table 2). Two groups did not receive any vaccination. Study 3 included four groups; unvaccinated, unchallenged (−), unvaccinated, challenged (+), empty vector vaccinated (pYD1 only), challenged, and pYD1-All 3 antigens vaccinated, challenged (Supplementary Table 2). All groups except one (negative control group) were challenged at day 22 with 4,000 *E. tenella* oocysts. The choice of dose level was based upon previous titration in this chicken line (21). All chickens were weighed and culled six days later. The caeca were examined for lesion scores as originally described by (22).

#### Study 4 (High dose challenge-broiler chickens)

A high parasite dose challenge study was then used to assess vaccine efficacy in broiler lines; as for the work in layers, pathological and performance parameters were evaluated. One hundred and fifty (150) mixed sex Cobb500 broiler day of hatch chicks were initially housed together on fresh litter. At day 7 chickens were weighed, wing-tagged and divided into four groups of 35 chicks, each group was then housed in separate pens with control groups (unvaccinated/unchallenged (−), and unvaccinated/challenged (+)) housed in separate rooms. Two groups received an oral yeast vaccine every 3-4 days from day 7 of age (five doses per chicken in total); the treatments were empty vector (pYD1 only) or a mixture of pYD1-EtAMA1, pYD1-EtIMP1 and pYD1-EtMIC3 (Supplementary Table 2). Due to an outbreak of colibacillosis within all groups, confirmed by bacterial culture of liver samples obtained post-mortem, all chickens were treated with enrofloxacin (Baytril®, Bayer, Leverkusen, Germany) at 10 mg Kg^−1^ for 3 days from days 16 to 18. All chickens were weighed at day 21 then all infected groups except one were challenged by oral inoculation with 15, 000 *E. tenella* oocysts. The choice of dose level was based upon previous studies with Cobb500 chickens (23), where a dose higher than that used with the Hy-Line chickens was required to achieve a comparable level of pathology. The unvaccinated/unchallenged negative control group received a mock challenge using PBS. From each group, a randomly selected cohort of 8-10 chickens were weighed and culled six days later to assess the pathological consequences of infection. The caeca from these chickens were examined for lesion scores as originally described by (22). At 10 days post infection the remaining chickens (n=19-21 / group) were weighed and culled. Feeders were emptied at time of challenge and food intake was then measured until chickens were culled to calculate feed conversion ratio (FCR) for each group by dividing total food consumed by total body weight gain (Day 21-31).

### 2.8 Isolation of total genomic DNA from caecal tissue for quantification of parasite replication

Genomic DNA (gDNA) was extracted from caecal tissues stored in RNAlater™ (Thermofisher Scientific) from Study 1. Caeca were homogenised in Buffer ATL using a Tissue ruptor homogenizer (QIAGEN) and then digested overnight at 56 °C in Buffer ATL and proteinase K, prior to extraction using the DNeasy Blood and Tissue DNA Kit (QIAGEN) according to manufacturer’s instructions.

### 2.9 Quantitative PCR for parasite replication

Quantitative PCR for assessment of *E. tenella* genome copy number in the caeca was performed as previously described to quantify parasite replication (24). Briefly, gDNA purified from caecal tissue was used as template for qPCR targeting *E. tenella* (RAPD-SCAR marker Tn-E03-116, primers F:TCGTCTTTGGCTGGCTATTC, R: CAGAGAGTCGCCGTCACAGT (25)) and chicken (tata-binding protein (TBP), primers F: TAGCCCGATGATGCCGTAT, R: GTTCCCTGTGTCGCTTGC (26)) genomes. Quantitative PCR was performed in 20 μL reactions in triplicate containing 10 μL 2 × SsoFast EvaGreen Supermix (Bio-Rad,), 1 μL of primers (3 μM F and 3 μM R), 8 μL of molecular biological grade water (Invitrogen) and 1 μL of gDNA or water as a negative no template control. Hard-shelled 96-well reaction plates (Bio-Rad) were sealed with adhesive film (Bio-Rad) and loaded into a Bio-Rad CFX qPCR cycler. Reactions were heated to 95°C for 2 min, prior to 40 cycles consisting of 95°C for 15 s then 60°C for 30 s with a fluorescence reading taken after each cycle. Melting curve analysis was performed consisting of 15 s at 95°C, before cooling to 65°C for 60 s, then heating to 95°C in 0.5°C increments for 0.5 s. Absolute quantification was performed against a standard curve generated using serially diluted plasmid DNA containing the amplicon of interest (EtenSCAR or ChickenTBP), to generate a standard curve ranging from 10^6^ copies to 10^1^ genome copies per mL. Parasite genome copy number was normalised by division with host (chicken) genome copy number.

### 2.10 Statistical analysis

Statistical analysis was carried out using GraphPad Prism 8 (Graph Software, LLC). One-way ANOVA was used to compare means of different groups for weight gain and parasite replication, D’Agostino-Pearson normality testing was performed to confirm a Gaussian distribution. The Kruskal-Wallis test was used to compare ranked means of challenge groups for lesion scores. The post-hoc multiple comparison test used for all parameters was Tukeys, and Spearmann rank correlation was used to assess correlations between parameters.

## 3 Results

### 3.1 Expression of E. tenella antigens in killed S.cerevisiae

Confirmation of protein expression by *S. cerevisiae* 24 h post-induction was provided by antibody staining and flow cytometry, indicating inducible expression of the *E. tenella* antigens EtAMA1, EtIMP1 or EtMIC3 based upon detection of the 3’ V5 epitope tag (examples shown in Supplementary Figure 1). Successful killing of each *S. cerevisiae* vaccine line was confirmed by the absence of growth on (i) minimal dextrose plates supplemented with 1 % leucine and 2 % glucose, and (ii) YNB-CAA medium containing 2 % glucose, after five days incubation. Heat killing at 56 °C for 1 h as per study 1 was found to be less consistent in heat inactivating yeast with occasional growth observed compared to the higher temperature of 95 °C for 2 min, as per study 2. Thus, the latter treatment was used for subsequent studies.

### 3.2 Significant reduction in parasite replication following oral yeast vaccination and low dose challenge (Study 1)

A significant decrease in parasite genome copy number at 5 days post-infection was observed in chickens vaccinated orally with *S. cerevisiae* expressing either EtAMA1Cit or EtIMP1Cit alone, and for those given an admixture of *S. cerevisiae* expressing both antigens, compared with unvaccinated, challenged chickens (p<0.01) (Figure 1). The use of a mixture of *S.cerevisiae* expressing both antigens reduced parasite load significantly compared with *S.cerevisiae* expressing either EtAMA1Cit (p<0.05 or EtIMP1Cit alone (p<0.01), and compared with live oocyst vaccination (p<0.01) or vaccination using recombinant EtIMP1 protein (p<0.01). There was no significant difference in parasite load (number of genomes detected by qPCR) between chickens vaccinated with *S.cerevisiae* expressing either EtAMA1Cit or EtIMP1Cit alone and those given live oocyst vaccination(p>0.05). Control groups performed as anticipated; there was no significant difference between chickens vaccinated with *S. cerevisiae* containing the empty pYD1 vector (p>0.05) compared with unvaccinated, challenged chickens. Chickens vaccinated with a live oocyst dose at day 7 or vaccinated with recombinant EtIMP1 protein showed a significant decrease in parasite genome copy number compared with unvaccinated, challenged chickens (p<0.01). Mean percentage reduction in parasite load, compared with unvaccinated, challenged chickens, for chickens vaccinated with *S. cerevisiae* expressing either EtAMA1 or EtIMP1 (64.7 % and 54.7 %, respectively) was comparable with vaccination with recombinant EtIMP1 protein given intramuscularly (59.8 %). Whilst combined vaccination with *S. cerevisiae* expressing EtAMA1 and EtIMP1 reduced mean parasite load further to 86.2 %.

**Figure 1.**
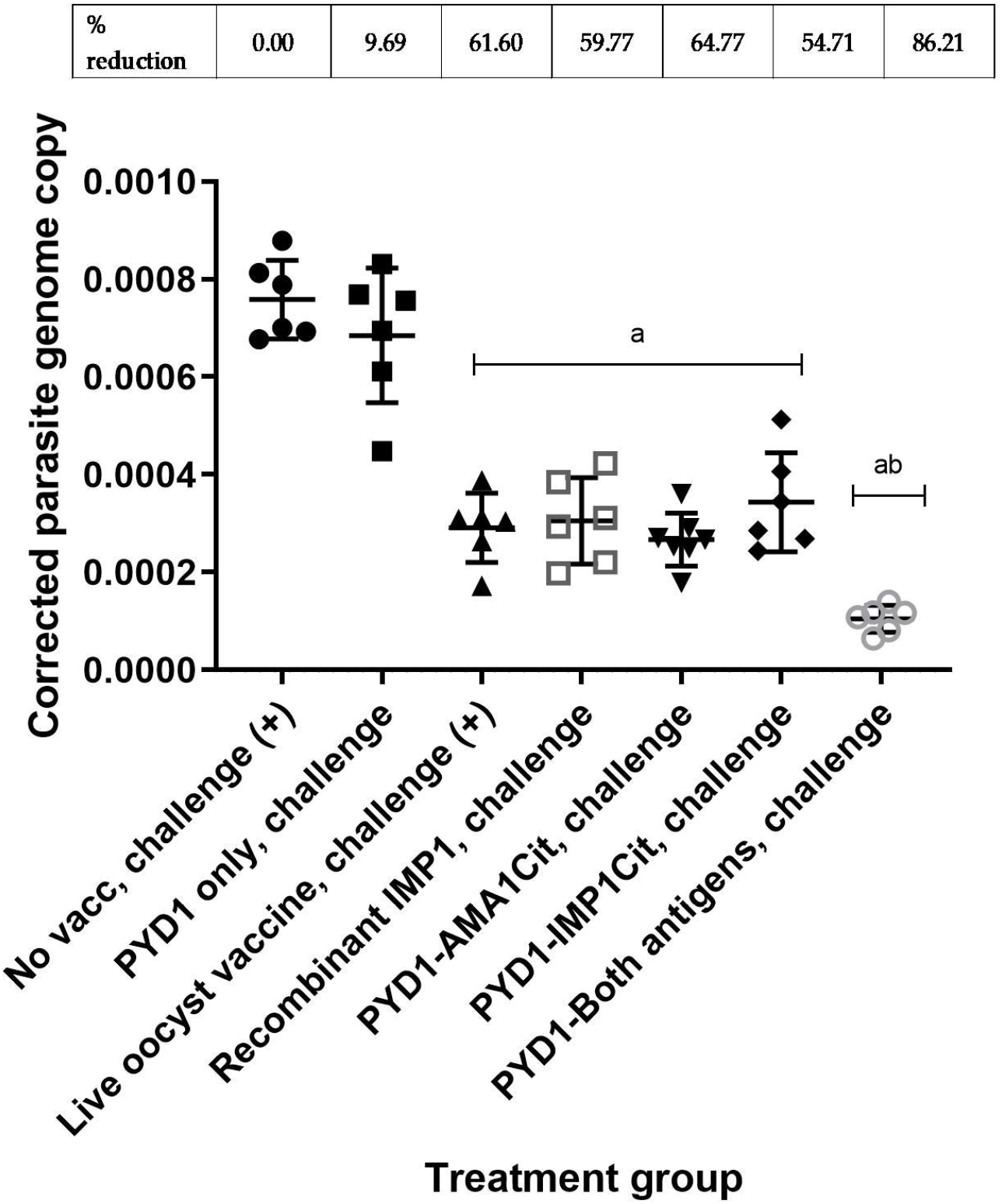
*Eimeria tenella* replication assessed by qPCR of genomic DNA extracted from caeca following low dose challenge in Hy-line brown layer chickens (Study 1). Treatment groups are shown on the x-axis and *E. tenella* genome copy number corrected using chicken TBP copy number is on the y-axis. Each marker represents one chicken (n= 6-8 per group). Mean and standard deviation for each group is shown. Groups with significantly different mean corrected parasite copy number compared with unvaccinated, challenged chicken group are shown with letter a. The group with significantly mean corrected parasite copy number compared with all other groups is shown with letters ab. Percentage reduction in mean corrected parasite copy number compared with unvaccinated, challenged chicken group are shown above graph.

### 3.3 Caecal lesion scores reduced in a proportion of layer chickens vaccinated with combination of all three antigens expressed in S. cerevisiae in high challenge study (Studies 2 and 3)

Although there was no statistical difference in mean lesion score between the vaccinated chickens and unvaccinated chickens in Study 2 (Figure 2A), it was apparent that a proportion of chickens that received a mixture of *S. cerevisiae* expressing EtAMA1, EtIMP1 and EtMIC3 had either no visible caecal lesions (5/13) or a lesion score of 1 (3/13). This reduction in lesion score in some chickens was less marked in those which received *S. cerevisiae* expressing only one of the three antigens. In study 3, which studied are larger group of chickens but otherwise followed a comparable study design, a statistically significant reduction in caecal lesion score was observed in groups of chickens vaccinated using a mixture of *S. cerevisiae* expressing EtAMA1, EtIMP1 and EtMIC3 compared with unvaccinated, challenged chickens (p<0.01) (Figure 2B). As in study 2, there was marked variability in lesion scores between individual vaccinated chickens (range 0-3).

**Figure 2.**
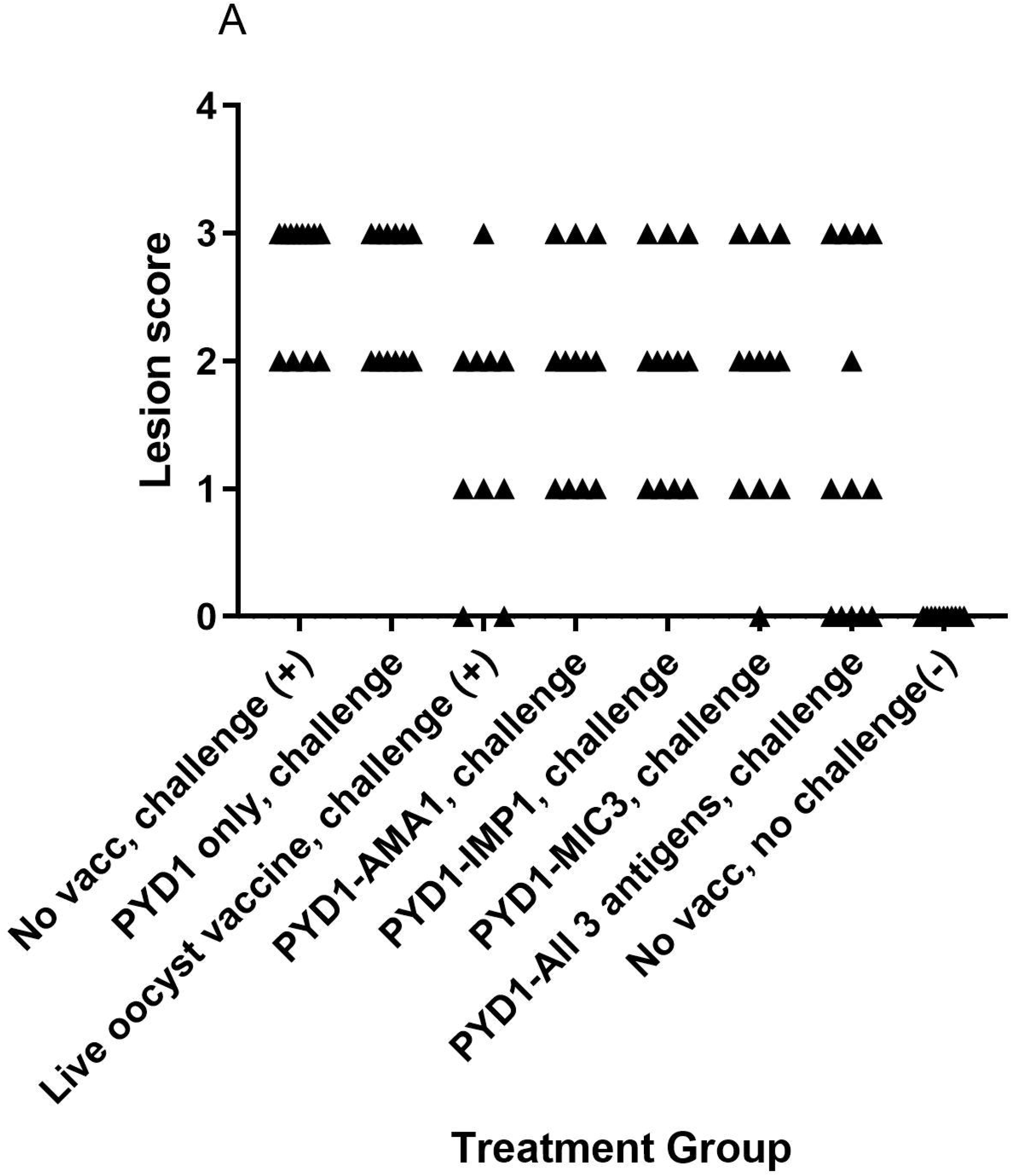
Caecal lesion scores six days post high dose *E. tenella* infection in Hy-line brown layer chickens (Studies 2 and 3) Treatment groups are shown on the x-axis and lesion scores are shown on the y-axis. Each marker represents one chicken (Study 2: n=10-13, Study 3: n=33-34). A. Caecal lesion scores for study 2. B. Caecal lesion scores for study 3. Groups with significantly different mean lesion score compared with the equivalent unvaccinated, challenged chicken group are shown with an asterisk (*).

### 3.4 Pre and post E. tenella challenge weight gain in layer chickens was unchanged by E. tenella infection after low and high dose challenge (Studies 1, 2 and 3)

No significant difference was noted in weight gain between unvaccinated, unchallenged and unvaccinated, challenged Hy-Line layer chickens in studies 1 and 2. Average weight gain in the six days post-challenge was 106.7 g ± 14.02 g and 85.18g ± 7.125 g in the unvaccinated, unchallenged groups, and 97.17 g ± 15.74 g and 90.67 g ± 11.19 g in the unvaccinated, challenged groups (studies 1 and 2, respectively). In the absence of a significant difference between positive and negative controls in these studies, weight gain was not assessed as a performance parameter. A statistical difference in weight gain post-challenge was observed in study 3 with the unvaccinated, unchallenged group (78.12 g ± 11.03 g) demonstrating higher body weight gain compared with the unvaccinated, challenged group (65.58 g ± 16.02 g; p<0.01). There was no significant improvement in weight gain post-challenge in *S. cerevisiae* vaccinated chickens compared with unvaccinated, challenged controls in study 3 (65.35 g± 15.35 g; p>0.05). Chickens used in these studies were commercial layer chickens and there was no anticipated impact on body weight gain in the short time period (5-6 days) studied post-challenge.

### 3.5 Caecal lesion scores were reduced in a proportion of broiler chickens vaccinated with a combination of all three S. cerevisiae expressed antigens (Study 4)

As described for the layer chickens, a wider range in lesion score was observed in broiler chickens vaccinated with a mixture of *S. cerevisiae* expressing all three antigens compared to those left unvaccinated (unvaccinated, challenged: average 3.3, range 3-4; test vaccinated, challenged: 2.6, 2-3; Figure 3). However, there was no statistical difference in mean lesion score at day 6 post-challenge (Figure 3).

**Figure 3.**
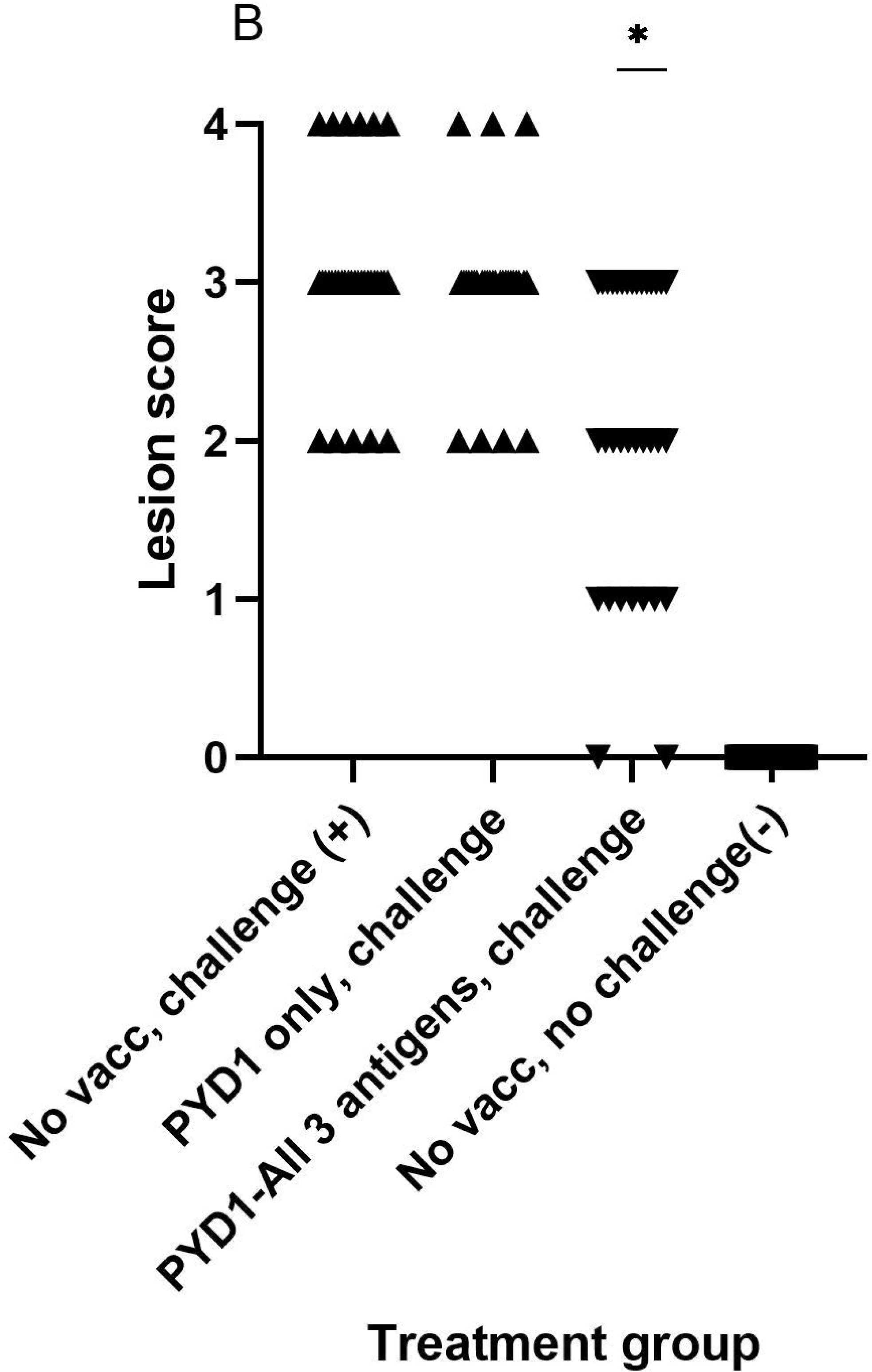
Caecal lesion scores six days post high dose *E. tenella* challenge in Cobb500 broiler chickens (Study 4). Treatment groups are shown on the x-axis and lesion scores are shown on the y-axis. Each marker represents one chicken (n=8-10).

### 3.6 Significant increase in body weight gain post E. tenella challenge and improved food conversion ratio following oral yeast vaccination in broiler chickens

A significant increase in body weight gain was observed for Cobb500 chickens vaccinated orally with a mixture of *S. cerevisiae* expressing all three antigens compared with unvaccinated, challenged chickens over the ten days following high level *E. tenella* challenge (p < 0.01; Figure 4). There was also a significant increase in body weight gain in vaccinated chickens compared to mock vaccinated chickens that received the empty pYD1 vector (p<0.05). There was a significant difference in body weight gain between unvaccinated, unchallenged chickens and unvaccinated, challenged chickens (p<0.05). When chickens were grouped by sex the significant increase in body weight gain in vaccinated compared with unvaccinated, challenged chickens (study days D21-D31) remained; the increase in body weight was more significant in females (p < 0.001) than males (p<0.05). There was no significant difference in body weight gain between groups pre-challenge (D7-21) (p>0.05). There was no significant difference in mean body weight between groups at any of the time points evaluated (D7, D21, D31) (p>0.05) (Table 1).

**Figure 4.**
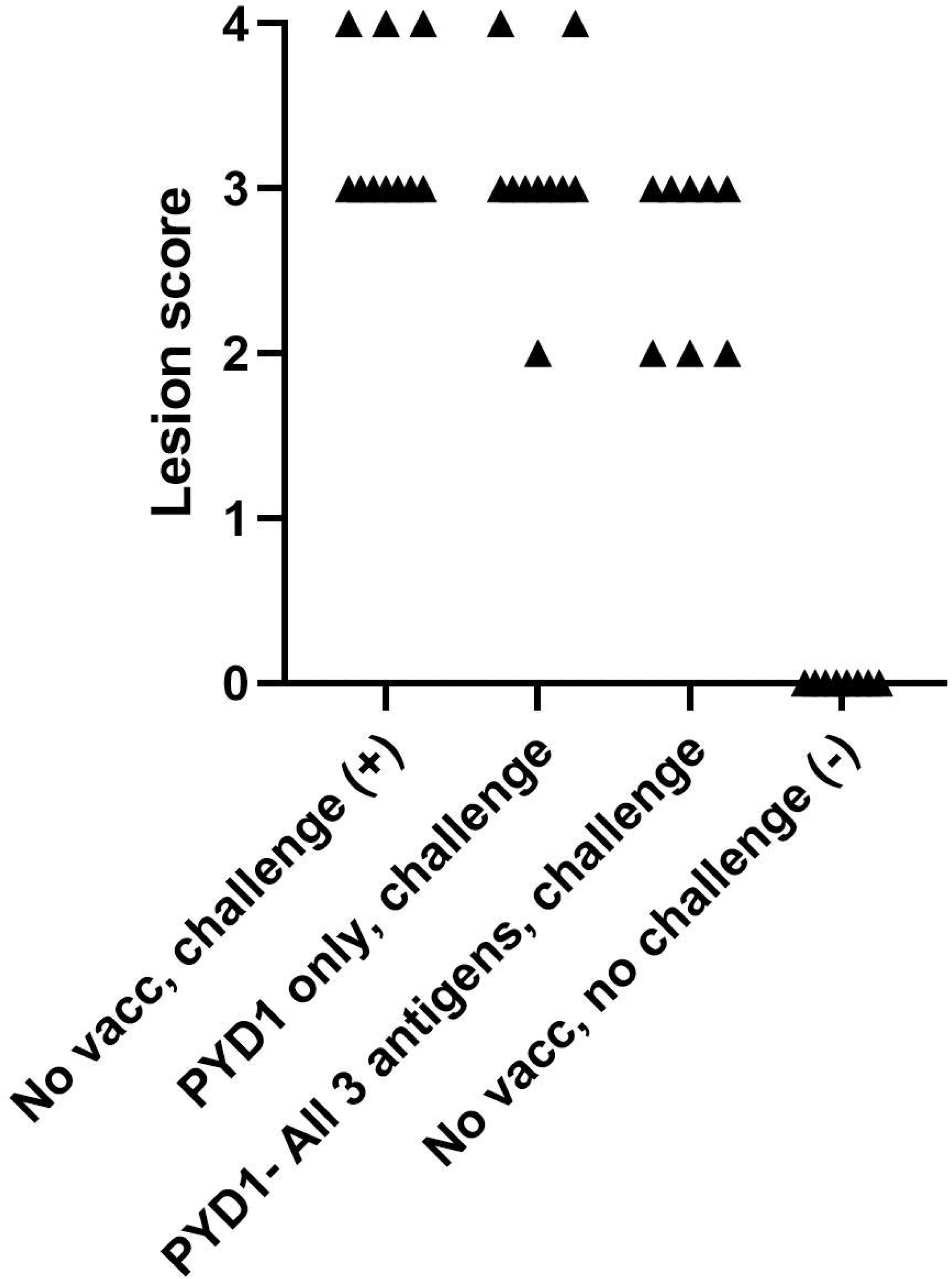
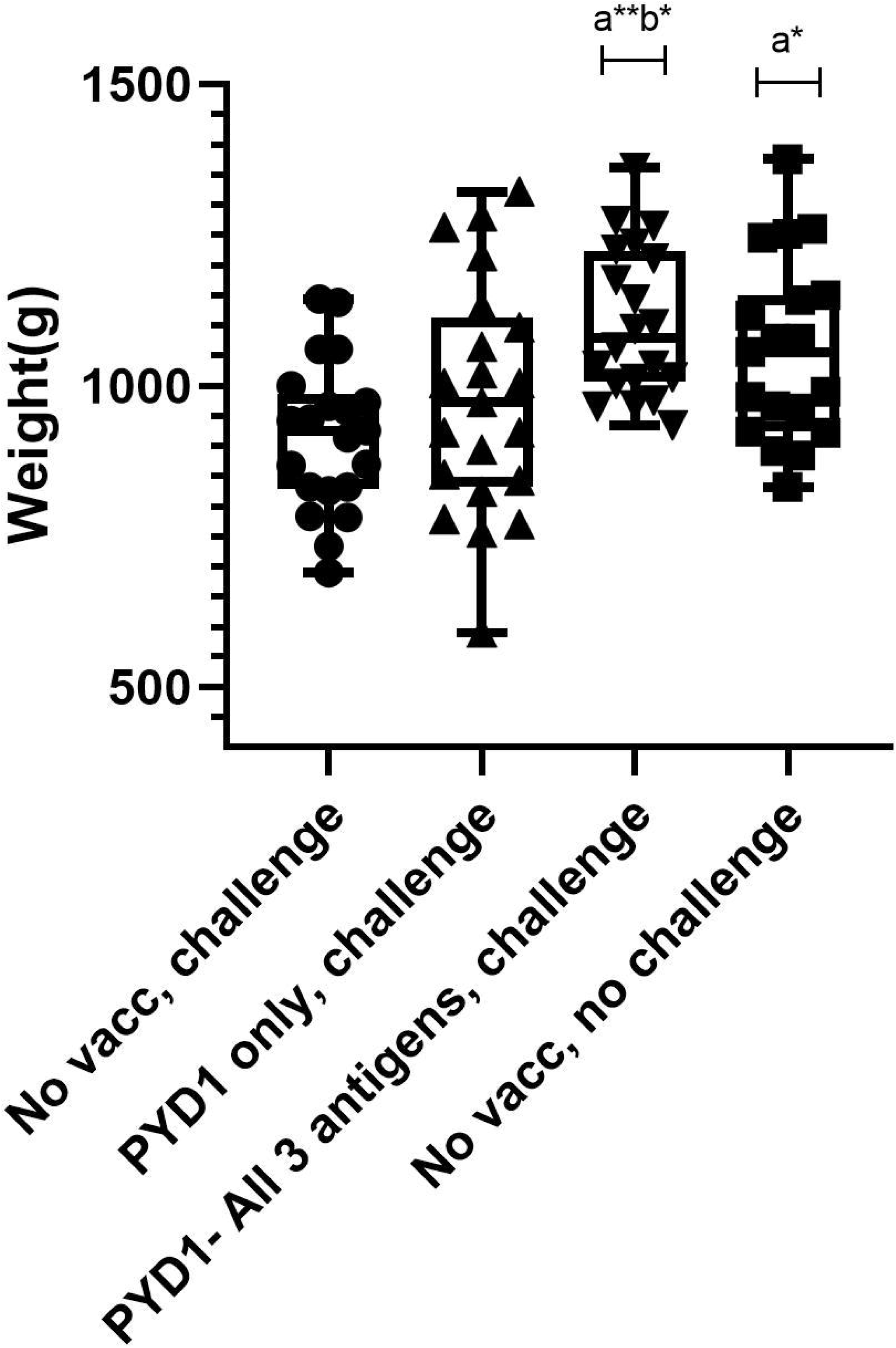
Body weight gain post high dose *E. tenella* challenge in Cobb500 broiler chickens (Study 4). Treatment groups are shown on the x-axis and body weight gain in grams from Day 21-31 is shown on the y-axis. Each marker represents one chicken (n= 19-21 per group). Groups with significantly different body weight gain compared with the unvaccinated, challenged chicken group are denoted by the letter a and those significantly different from empty vector (pYD1 only) vaccinated challenge group denoted by the letter b. One asterisk (*) denotes significance level p<0.05, two asterisk (**) denotes significance level p<0.01 (One way ANOVA, Tukey multiple comparison correction).

**Table 1.** Summary of Cobb500 broiler chicken body weight in the high dose *E .tenella* challenge study (Study 4). Chickens were weighed at day 7 (pre-vaccination), day 21 (day of challenge) and day 31 (10 days post-challenge).

| Group | Mean body weight(g), (SD) |  |  | Mean body weight gain(g) |  |
| --- | --- | --- | --- | --- | --- |
|  | Day 7 | Day 21 | Day 31 | Pre-challenge(D7-21) | Post-challenge(D21-31) |
| Unvaccinated, challenged (+) | 212<br>(13.58) | 1139(112.<br>2) | 2036<br>(191.3) | 926.9 (103.9) | 916.3 (123.5) |
| Unvaccinated, unchallenged (-) | 216.1<br>(15.23) | 1149<br>(124.2) | 2172<br>(263.3) | 933 (114.7) | 1058 (150.6)<sup>a</sup> |
| Empty vector vaccinated(pYD1 only),<br>challenged | 208.3<br>(18.38) | 1112(107.<br>2) | 2094<br>(300) | 903.8 (100.9) | 977.1 (193.4) |
| pYD1-All 3 antigens, challenged | 211.6<br>(15.05) | 1108<br>(116.1) | 2223<br>(216.2) | 896.8 (108.4) | 1105 (124.1)<sup>ab</sup> |
| Number chickens per group | 28-32 | 28-32 | 19-21 | 28-32 | 19-21 |
| F statistic, p value (ANOVA) | 1.261, 0.29 | 0.8866,<br>0.45 | 2.319,<br>0.08 | 0.7825, 0.51 | **6.317, 0.0007** |

<sup>a</sup>-significant difference compared to unvaccinated, challenged group, <sup>b</sup>-significant difference compared to empty vector vaccinated (pYD1 only), challenged group

Food conversion ratio (FCR) was calculated for each group, together with total body weight gain of chickens culled at six- and ten-days post-challenge (Table 2). FCR was lowest in chickens vaccinated with a mixture of *S. cerevisiae* expressing all three antigens at 1.52, comparable to unvaccinated, unchallenged chickens with an FCR of 1.56. Chickens vaccinated with *S. cerevisiae* with empty pYD1-vector had a higher FCR of 1.65 comparable to unvaccinated, challenged chickens with a FCR of 1.67. Statistical comparison of the differences in FCR between groups was not possible because values were calculated per treatment group, rather than for individual chickens.

**Table 2.** Summary of food conversion ratio (FCR) in Cobb500 broiler chickens in the high dose *E. tenella* challenge study (Study 4). FCR was calculated together with total body weight gain of chickens culled at six and ten days post-challenge.

| Group | Total weight gain post-challenge (kg) D21-D31 | Total food consumed(kg)D21-D31 | Food conversion ratio |
| --- | --- | --- | --- |
| Unvaccinated, challenged(+) | 25.171 | 42.14 | 1.67414882 |
| Unvaccinated, unchallenge(-) | 25.174 | 39.23 | 1.55835386 |
| Empty vector vaccinated (pYD1 only), challenged | 26.653 | 44.00 | 1.65084606 |
| pYD1-All 3 antigens, challenged | 26.177 | 39.78 | 1.51965466 |

## Discussion

Development of novel recombinant or subunit vaccines against *Eimeria* species in chickens has been limited thus far. Despite many promising pilot studies with various antigens, none have progressed to commercial products (27). The absence of an efficacious, cost-effective and scalable system for routine vaccination of broilers remains a persistent challenge for *Eimeria,* as well as other pathogens. In this study we sought to address some of these barriers to commercialisation for *Eimeria* vaccines by developing an oral inactivated whole yeast-based vaccine which could be produced easily and cheaply, and potentially be administered in-feed. Taking a panel of candidate immunoprotective antigens validated previously using recombinant protein and/or DNA vaccination screens (4–6), we have demonstrated here that *S. cerevisiae* yeast expressing *E. tenella* antigens could be produced and delivered as a whole inactivated yeast vaccine safely to layer and broiler chickens. Further, such vaccines can be effective in reducing *E. tenella* replication in the caeca, reduce intestinal lesion score, and improve body weight gain and food conversion post-challenge. *Saccharomyces cerevisiae* expressing *E. tenella* antigens were heat killed and freeze-dried before use as a whole yeast vaccine, thus they were no longer classified as genetically modified organisms (GMO) which simplifies future licensing. Heat-killing of *S. cerevisiae* has been described elsewhere(15, 28), although there is little published data on validating methods of heat-killing for yeast and specification by national/international regulators will likely be needed. From this study, it appeared that heat killing at high temperature (95°C) for two minutes was more reliable than a longer incubation at lower temperature, but this may vary depending on the concentration of yeast particles incubated and method of heating. Previous studies have demonstrated that protein antigen stability in yeast can be maintained for up to a year even at room temperature, making this system ideally suited for use in developing countries where cold-chain access may be limited (29). Based on previous studies heat-killing does not appear to impact immunogenicity (reviewed in (30)), although this was not within the scope of our study.

Vaccination with *S. cerevisiae* expressing EtAMA1 and EtIMP1, alone or together, was successful in reducing parasite replication following low level *E. tenella* challenge. Parasite replication was assessed in the context of a low level parasite challenge since the *Eimeria* crowding effect(19) can be expected to obscure partial protective responses at higher levels of challenge, as illustrated in a recent dose titration study using layer-breed chickens(21). Yeast vaccination with a single antigen was comparable to vaccination by injection with the equivalent recombinant EtIMP1 protein in the low dose challenge. Previous studies have demonstrated the efficacy of these antigens when delivered subcutaneously or intramuscularly as protein or DNA vaccines (4–6). More recent studies have delivered these antigens orally (31, 32). EtAMA1 and EtIMP1 expressed and delivered in a live recombinant bacterial *Lactococcus lactis* vaccine resulted in reductions in oocyst output and lesion score (31, 32). Similarly, EtAMA1 and EtMIC2 co-expressed in *Lactobacillus plantarum* and delivered as an oral vaccine reduced oocyst output and lesion score following challenge (33). As noted for most recombinant antigen-based vaccines for *Eimeria* (reviewed in (27)), our yeast-based vaccine reduced parasite replication but did not completely prevent parasite development and shedding. Modern ionophore anticoccidial formulations reduce replication of drug susceptible *Eimeria* populations by 82-97% (34) and it is likely that a successful recombinant vaccine will need to achieve a comparable reduction. It is noteworthy that vaccination using a mixture of yeast expressing EtAMA1 with EtIMP1 achieved 86.2% reduction in parasite replication, which is in the range of commercially viable levels of efficacy. Moreover, in a field situation, reduction in parasite replication and oocyst output is likely to be boosted by low levels of parasite escape and recycling as seen with current live and live-attenuated vaccines, as well as with the use of current anticoccidial drugs (3). Furthermore, it will likely be possible to improve vaccine efficacy by altering doses or treatment schedules, which will be examined in future studies.

Assessment of the effects of *Eimeria* infection on enteric pathology and production parameters require high levels of parasite challenge, carefully titrated to achieve a measurable phenotype without mortality (21). Performance parameters such as body weight gain and FCR can most usefully be assessed in broiler-type chickens that have rapid and efficient growth (35). Slower growing layer-type chickens are less likely to provide discriminatory phenotypes in a short study, as seen here using Hy-Line Brown layer chickens. The results of our high challenge broiler study indicate significant improvement in key production traits of body weight gain and food conversion rates in chickens vaccinated with *S. cerevisiae* expressing *E. tenella* antigens, suggesting that disease was sufficiently reduced to allow for sustained body weight gain required by the broiler industry. Indeed, average body weight gain post-challenge in *S. cerevisiae- E. tenella* vaccinated broilers was higher than in unchallenged chickens, better even than the 90% body weight maintenance observed with monensin treated chickens reported previously (34). This difference might reflect compensatory growth, or a positive additive effect of dietary supplementation using *S. cerevisiae* (10, 11). Reducing the consequences of *Eimeria* infection can also be expected to improve chicken welfare. Further field studies are required to evaluate the use of this vaccine under commercial broiler conditions to confirm its applicability to the market.

While yeast-vectored anticoccidial vaccination improved measures of parasite replication and broiler performance, the impact of vaccination on lesion score was more nuanced. Considerable inter-animal variation was observed in lesion scores in vaccinated chickens, with some reductions statistically significant while others were not. Lesion scores were also subjectively higher within the broiler study compared to the high challenge layer studies, reflecting differences in genetic resistance/susceptibility between these chicken types. Development of lesions following *Eimeria* infection is complex and is likely a combination of host genetics, level of parasite damage to epithelial cells and host inflammatory response (36). Genome-wide association studies (GWAS) have suggested a significant host genetic influence on the outcome of secondary *Eimeria* infection (37), a feature that might also apply to the success of vaccination. It has been suggested that lesions deriving from primary infection might suggest severe disease, whilst lesions in chickens vaccinated with live or live attenuated vaccines arising post vaccination or following subsequent challenge are not necessarily indicative of a lack of protection from disease (35, 38). Previous studies have demonstrated the presence of lesions in chickens vaccinated with live attenuated vaccines, however lesions were less severe in vaccinated chickens and not always associated with presence of endogenous parasites compared to unvaccinated chickens where parasite numbers were high (39). Although our study did not microscopically examine caecal lesions, evidence of reduced parasite replication in the caeca as demonstrated by the qPCR data could support a similar phenomenon following vaccination with our yeast vaccines.

In addition to antigen expression and delivery, a killed yeast vaccine can also provide an immunostimulatory adjuvating effect. It is well established that *S. cerevisiae* yeast are immunogenic and can be taken up and activate macrophages and dendritic cells (40) through receptors such as the mannose receptor and Dectin-1, which also have been shown to be expressed on mammalian M-cells (41). Whilst this mechanism makes them ideal for generating an antigen-specific adaptive immune response following antigen presentation through MHC, they also stimulate an innate immune response. Indeed, studies feeding yeast cell-wall components to chickens infected with *Eimeria* have demonstrated reduction in parasite replication and improvement in production traits (9–11). In the broiler study (Study 4), the improvement in body weight gain observed post-challenge appeared to be independent of the “yeast effect” with a significant increase in chickens vaccinated with *S. cerevisiae* expressing *E. tenella* antigens compared with those given *S. cerevisiae* empty vector control. Nonetheless, the beneficial effects of dietary yeast supplementation can add value to a vectored anticoccidial vaccine; dose optimisation will likely be required.

Any novel *Eimeria* vaccine will need to incorporate antigens that stimulate immune responses protective against more than one *Eimeria* species for it be a viable alternative to anticoccidial drugs or existing live and live-attenuated vaccines. It is well established that there is little to no cross immune protection against heterologous challenge between *Eimeria* species, indicating a requirement for additional antigens(42). *Eimeria tenella* was selected for initial proof of concept being both well described with established infection models and also an important species in terms of prevalence and pathogenicity (17, 43). *Eimeria tenella* is also recognised as one of the least immunogenic of the *Eimeria* that infect chickens (18), which suggests that protection achieved here could be improved when using equivalent antigens derived from other, more immunogenic, species. All current live anticoccidial vaccines target *Eimeria acervulina* and *E. maxima*, in addition to *E. tenella*, as a core unit (44). Some vaccines formulated for broiler chicken markets have established a more specific identity by inclusion of other species such as *E. mitis* (e.g. Paracox-5 or HuveGuard MMAT) as well as *E. praecox* (e.g. Evant). Species such as *E. brunetti* and *E. necatrix* are usually only required in vaccines for longer lived layer or breeder chickens (44). Future studies should focus on the addition of antigens from *Eimeria acervulina* and *E. maxima*, especially important in North America (45), which in combination with *E. tenella* would represent the three species most costly to global chicken production.

In conclusion we have demonstrated that a heat killed oral *S. cerevisiae* vaccine expressing *E. tenella* antigens is safe and effective in reducing parasite replication following challenge with *E. tenella*. Future work should extend examination of the impact of vaccination on production traits such as body weight gain and food conversion ratio in broiler chickens during challenge by *E. tenella* as well as other key *Eimeria* species to ensure this approach is a viable alternative to anticoccidial drugs.

## Author Contributions

Conceptualisation, D.B., D.W. and F.T.; methodology, F.S., D. B., M.N. and T.K..; validation, F.S. and D.B. formal analysis, F.S.; investigation, F.S., M.N. and T.K.; writing—original draft preparation, F.S.; writing—review and editing, D.B., D.W. and F.T.; visualisation, F.S.; supervision, D.B., D.W. and F.T.; project administration, F.S., E.A., V.M-H. and S.K ; funding acquisition, D.B., D.W. and F.T. All authors have read and agreed to the published version of the manuscript.

## Acknowledgments

The authors would like to thank Iván Pastor-Fernández, Kelisandia Aguiar-Martins, Gonzalo Sanchez Arsuaga, Eleanor Karp-Tatham and Michelle Jones for assistance with animal studies.

## Data availability statement

The data presented in this study are available on request from the corresponding author.

