## Supplementary material for "A novel whole yeast-based subunit oral vaccine against Eimeria tenella in chickens": Suppl. data

**Supplementary Data**

**Supplementary Figure 1:**

Confirmation of antigen expression by antibody detection of V5 tag on 3’ end of antigenic protein by flow cytometric detection of AF488 conjugated antibody. Fluorescence in yeast transfected with pYD1 expressing EtAMA1 is shown in **panel A**, for yeast transfected with pYD1 expressing EtIMP1 in **panel B** and for yeast transfected with pYD1 expressing EtMIC3 in **panel C**. Data representative of time point 0 h (before induction of expression) and 24 h after induction of expression are shown as scatter plots and histograms showing detection of AF.

A	0h	24h

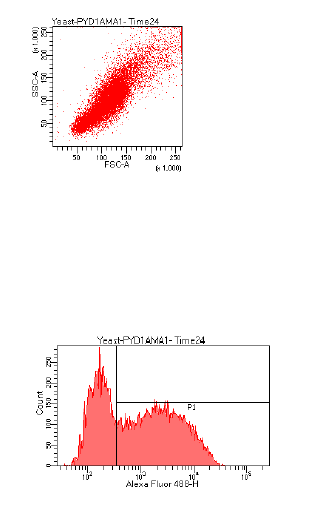

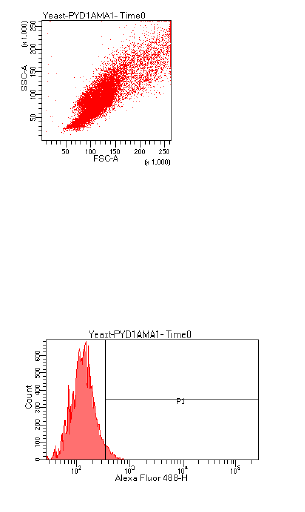

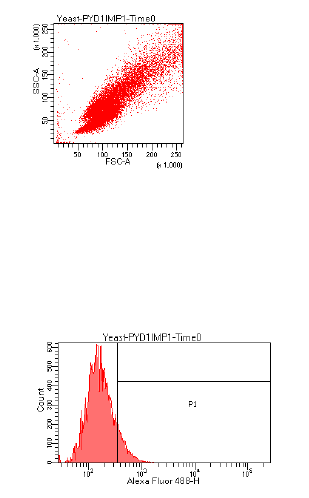

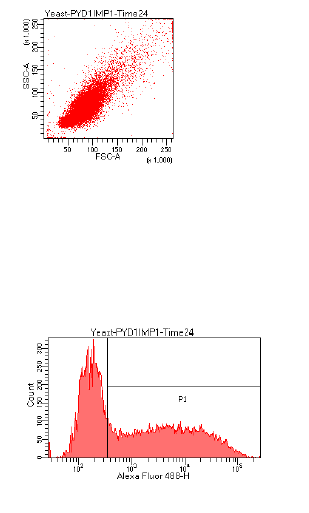

B	0h	24h

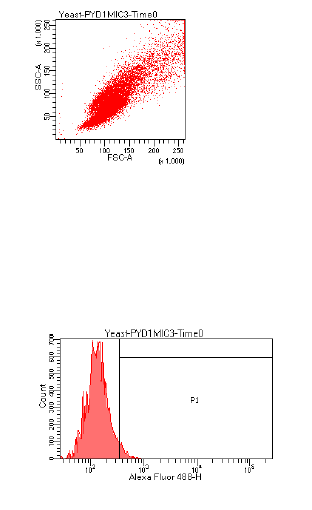

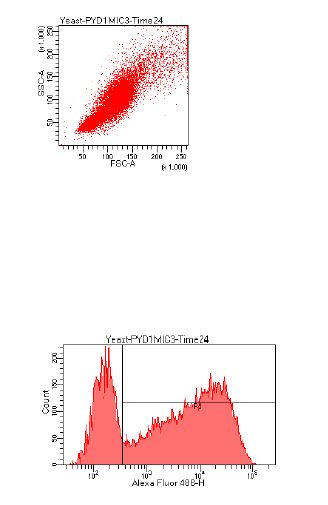

C	0h	24h

**Supplementary Table 1: Nutritional content of feed in Cobb500 broiler chicken study (Study 4)**

| **Nutrient** | **Starter** | **Grower** | **Finisher** |
| --- | --- | --- | --- |
| Oil EE | 6.3685 | 6.8778 | 7.1374 |
| Protein | 21.4436 | 20.2493 | 18.7742 |
| Fibre | 3.1325 | 3.0780 | 3.0290 |
| Ash | 6.0665 | 5.7290 | 5.5526 |
| ME-P | 12.8503 | 13.1191 | 13.3096 |
| Tlysine | 1.4343 | 1.2717 | 1.1619 |
| AvLysine | 1.3356 | 1.1794 | 1.0780 |
| Meth | 0.6900 | 0.6255 | 0.5555 |
| M+C | 1.0155 | 0.9353 | 0.8444 |
| Threo | 0.9013 | 0.8517 | 0.7861 |
| Trypt | 0.2523 | 0.2364 | 0.2157 |
| Calcium | 0.9755 | 0.9241 | 0.9111 |
| Phos | 0.7319 | 0.6600 | 0.6407 |
| AvPhos | 0.4848 | 0.4227 | 0.4163 |
| Salt | 0.3092 | 0.3077 | 0.3056 |
| Sodium | 0.1749 | 0.1751 | 0.1751 |
| Vit A | 13.5000 | 10.0000 | 10.0000 |
| Vit D3 | 5.0000 | 5.0000 | 5.0000 |
| Vit E | 100.0000 | 100.0000 | 100.0000 |
| **Raw Material** |  |  |  |
| Barley Raw Ground | 10.5000 | 8.4000 | 7.2000 |
| Wheat Raw Ground | 50.0000 | 55.0000 | 60.0000 |
| Soya Ext Hipro | 26.0000 | 23.0000 | 19.0000 |
| Full Fat Soya Masham | 5.0000 | 5.0000 | 5.0000 |
| L Lysine batch | 0.4000 | 0.3000 | 0.3000 |
| DL Methionine batch | 0.4000 | 0.3500 | 0.3000 |
| L Threonine batch | 0.1500 | 0.1500 | 0.1500 |
| Soya oil | 4.0000 | 4.5000 | 4.7500 |
| Limestone trucal | 1.2500 | 1.2500 | 1.2500 |
| Monocalcium phosphate | 1.5000 | 1.2500 | 1.2500 |
| Salt | 0.2500 | 0.2500 | 0.2500 |
| Sodium bicarbonate | 0.1500 | 0.1500 | 0.1500 |
| Br. Trial sta. pmx. batch | 0.4000 | 0.0000 | - |
| Br. Trial gro. pmx. batch | 0.0000 | 0.4000 | 0.4000 |

**Supplementary Table 2 Experimental study groups in each study**

| Study No. (description) | Group | No. of chickens |
| --- | --- | --- |
| 1 | Unvaccinated, unchallenged (-) | 6 |
| (low dose challenge-layers) | Unvaccinated, challenged(+) | 6 |
|  | Live oocyst vaccinated, challenged | 6 |
|  | Recombinant IMP1 vaccinated, challenged | 6 |
|  | Empty vector vaccinated(pYD1 only), challenged | 6 |
|  | pYD1-AMA1Cit vaccinated, challenged | 7 |
|  | pYD1-IMP1Cit vaccinated, challenged | 6 |
|  | pYD1- Both antigens vaccinated, challenged | 6 |
| 2 | Unvaccinated, unchallenged (-) | 11 |
| (high dose challenge-layers) | Unvaccinated, challenged(+) | 12 |
|  | Live oocyst vaccinated, challenged | 10 |
|  | Empty vector vaccinated(pYD1 only), challenged | 12 |
|  | pYD1-AMA1 vaccinated, challenged | 12 |
|  | pYD1-IMP1 vaccinated, challenged | 12 |
|  | pYD1-MIC3 vaccinated, challenged | 12 |
|  | pYD1- All 3 antigens vaccinated, challenged | 13 |
| 3 | Unvaccinated, unchallenged (-) | 33 |
| (high dose challenge-layers) | Unvaccinated, challenged(+) | 33 |
|  | Empty vector vaccinated(pYD1 only), challenged | 34 |
|  | pYD1- All 3 antigens vaccinated, challenged | 34 |
| 4 | Unvaccinated, unchallenged (-) | 28 |
| (high dose challenge-broilers) | Unvaccinated, challenged(+) | 31 |
|  | Empty vector vaccinated(pYD1 only), challenged | 32 |
|  | pYD1- All 3 antigens vaccinated, challenged | 28 |
